# Position-wise Fold-Switch Prediction in Metamorphic Proteins: A Comparison of Handcrafted and Language Model Features

**DOI:** 10.64898/2026.09.30.755750

**Authors:** Pranav Machingal, Nandyala Hemachandra, Petety V Balaji

## Abstract

In this work, we focus on predicting fold-switching phenomena observed in certain protein structures. Proteins that undergo fold-switching are called metamorphic proteins. We model fold-switch prediction in proteins as a position/residue-wise classification task. We use various features extracted from a protein’s sequence as well as structure to train the classifiers. We propose a novel class of handcrafted parametrised features that encode the structural context of a residue. We also use embeddings from existing protein language models (PLMs) to train the classifiers. We train binary and 3-class classifiers, with the 3-class classifiers providing granular information on the type of fold-switch. In our computations, higher classification performance is achieved for binary prediction as compared to 3-class granular prediction. The hand-crafted features perform comparably with the PLM embeddings on our standard test set.

A rigorous error analysis by us during the review period revealed a critical data-leakage vulnerability in the metamorphic protein test data. This analysis also revealed that the PLM-based classifier suffers sequence-memorisation bias and fails to generalise to novel fold-switching sequences. On the other hand, one of our domain-informed handcrafted feature based classifier detects fold-switching without relying on sequence memory.

## 1 Introduction

Proteins are the workhorses of a cell. A class of proteins called globular proteins assume a well-defined fold (loosely synonymous with 3D structure) in their native state. Anfinsen [3] showed that both the amino acid sequence and environmental conditions dictate the fold of a globular protein. In the last two decades, some globular proteins have been found to be capable of (partly) switching folds [6].

This class of fold-switching proteins are referred to as metamorphic proteins. The fold-switch may be brought about by changes in pH, redox potential, membrane association, etc. It is not clear if fold switching is a universal property of proteins or is restricted to certain sequences or sequence regions. It is practically impossible to perform wet-lab experiments to identify fold-switching in proteins, since there are millions of conditions to screen.

*In this work, we aim to predict which regions in a protein are fold-switching*. Particulary, we formulate this as a position-wise (or residue-wise) prediction task. The structural context of a residue, at a given position in the protein, is encoded into a fixed-dimensional representation. These feature vectors are then used to train classifiers that predict whether the residue at the given position will be involved in fold-switching. We evaluate and compare the effectiveness of handcrafted and automated protein language model (PLM) based feature engineering in this task.

The challenges in computational modelling of fold-switching proteins are highlighted in [11]. Molecular dynamics (MD) simulations have been used for predicting fold-switching [20]. However, MD simulations are not easily scalable. [11] also discusses a class of deep learning-based methods that primarily use pretrained structure-prediction algorithms such as AlphaFold2 [9] and strategically manipulate their input data to predict alternate structural conformations of a protein sequence. These methods fail to predict alternate conformations for a given input sequence if alternate conformations for homologous (evolutionarily related) sequences are not part of the training data. More recent methods [7,18] make a binary prediction (fold-switching or not) for a given input sequence, instead of predicting alternate conformations. [7] exploits the disagreement of various homology based secondary structure prediction algorithms to predict if a query a sequence is metamorphic or not. [18] looks at the structural diversity of proteins that are homologous to the query sequence to predict if it is metamorphic or not.

In contrast to these existing methods, which either predict the complete alternate conformation or a sequence-level binary prediction for fold-switching, in our work, we make position/residue-wise prediction of fold-switching. For this classification task, instead of a homology-based analysis, we propose a novel class of handcrafted parametrised features that encode the structural context of a residue. The motivation for designing these features is that their interpretability could provide insights into fold-switching phenomena. A recent paper [13] illustrated domain insights via interpretability of handcrafted features in the context of protein structure classification. We also use embeddings extracted from existing multi-modal PLMs for this task.

Crucially, the evaluation of metamorphic protein classifiers is highly susceptible to implicit data leakage. This is because a given metamorphic protein has two distinct structural conformations in the dataset with the same sequence. We demonstrate in our evaluations how this leads to sequence memorisation by the classifiers. In particular, we observe sequence memorisation in PLM-based classifiers when analysing strictly leak-proof data. In contrast, we do not see this sequence memorisation in the best handcrafted feature based classifier. This highlights the need for rigorous benchmarking and domain-informed feature engineering in structural bioinformatics.

## 2 Background: Proteins

We provide background on the biological concepts discussed in this paper [8,14] that are relevant to our fold-switching prediction task.

### Globular proteins

Proteins can be globular, fibrous, intrinsically disordered, or functional amyloids. Among these, globular proteins are known to switch folds. In this paper, these are simply referred to as *proteins* for brevity.

### Amino acids

These are the building blocks of proteins. In general, proteins consist of 20 different types of amino acids, in various compositions. An amino acid is referred to as a ***residue*** when it connects with another amino acid via a peptide bond to form a protein.

### Protein sequence

The amino acids in a protein are connected to each other in a sequence via peptide bonds to form a linear chain. The sequence in which amino acids are linked is also referred to as the protein’s primary structure or simply the amino acid sequence.

### Secondary structure

In 3-dimensions, local contiguous segments of the protein sequence often fold into well-defined structures or otherwise have a random disordered structure. This local structure of a contiguous segment in a protein sequence is referred to as the secondary structure of that segment. The secondary structures are broadly considered to be of three types: *α*-helix, *β*-strand and coil (*c*; random/disordered). We refer to these simply as *α, β* and *c* respectively in the paper.

### Tertiary structure

It is the overall 3-dimensional structure of the protein defined by interactions between the amino acids, which may be close or distant in the sequence, and also defined by non-covalent interactions with the solvent molecules. The folded protein structure can be considered an origami of a series of connected secondary structure elements of varying lengths.

### Metamorphic proteins [6]

The proteins that can reversibly switch between at least two different stable protein structures are called metamorphic proteins. The fold-switch could be triggered by various biochemical factors. These proteins span the animal kingdom and biological functions [6]. The entire protein need not switch its fold. In general, the “fold-switching” in the structure of a metamorphic protein is characterised by the switching of one or more of its secondary structure elements from one type to another type.

### Monomorphic proteins

The proteins that are considered to have only one stable structure are called monomorphic proteins.

## 3 Problem formulation: fold-switching prediction as a classification task

The identification of metamorphic regions is formulated as a position-wise / residue-level classification task. Given an input protein structure, we aim to predict, for each of its residues, whether its secondary structure will change in an alternate conformation or not. If *S* is an input protein structure, then we want a feature engineering *ℱ* to get a fixed-dimensional feature vector *xi* for the residues at sequence position *i* of the structure, *xi* := *ℱ*(*S, i*). Then we want a classifier *h* : *xi ↦ yi* that predicts the structural switching at position *i*. For this, we look at two predictive paradigms,

- **Binary fold-switch prediction**: A binary classification task where the objective is to learn a mapping *hbin* : *xi ↦ yi* ∈ {0, 1}, where *yi* = 1 if a residue undergoes a secondary structure transition and otherwise, *yi* = 0.
- **3-class transition prediction**: A 3-class formulation aimed at predicting the exact target secondary structure state, *α*-helix/*β*-sheet/*c*oil, in the alternate conformation. The objective is to learn a mapping *hmulti* : *xi ↦ yi* ∈ {*α, β, c*}. This allows for the prediction of specific transition subtypes (e.g., *α* → *β, c* → *α*). If the secondary structure of the position *i* in the input structure is *si*, then the prediction is fold-switching if *yi* = *si*, else the prediction is that there is no fold-switching at position *i*.

This classification problem is highly imbalanced, as the number of fold-switching positions in a metamorphic protein is typically a smaller fraction of the protein’s total length.

## 4 Methodology

### 4.1 Dataset of metamorphic and monomorphic protein structures

For training and testing the classifiers, we derive the dataset from [18], comprising 386 protein structures from the Protein Data Bank (PDB) [5]. Out of these, 94 pairs (188 PDB structures) are metamorphic. A metamorphic ‘pair’ here means two alternate structural conformations of the same protein. The remaining 198 structures are considered to be monomorphic. Additional curation details are provided in appendix Section A. We use 80% of the structures for training and 20% for testing the classifiers. A sequence position *i* in a given structure can have 6 possible secondary structure fold-switches, {*si* → *s*^*′*^_*i*_|*si, s*^*′*^_*i*_ ∈ {*α, β, c*}, *si s*^*′*^_*i*_}. For positions with no fold-switch, *si* = *s*^*′*^_*i*_. To ensure that the fold-switch distribution across positions is similar between the train and test structures, we sample 10000 random 80:20 splits and choose the split with the minimum Jensen-Shannon divergence [15] between the empirical fold-switch distributions of the train and test data. Please see Fig. 1. From the training data structures, we use 10% of the sequence positions as a validation set to tune the classifier’s hyper-parameters. Thus, the train+val / test data split is structure-wise (80:20), while the train/val data split is position-wise (72:8).

**Fig 1:**
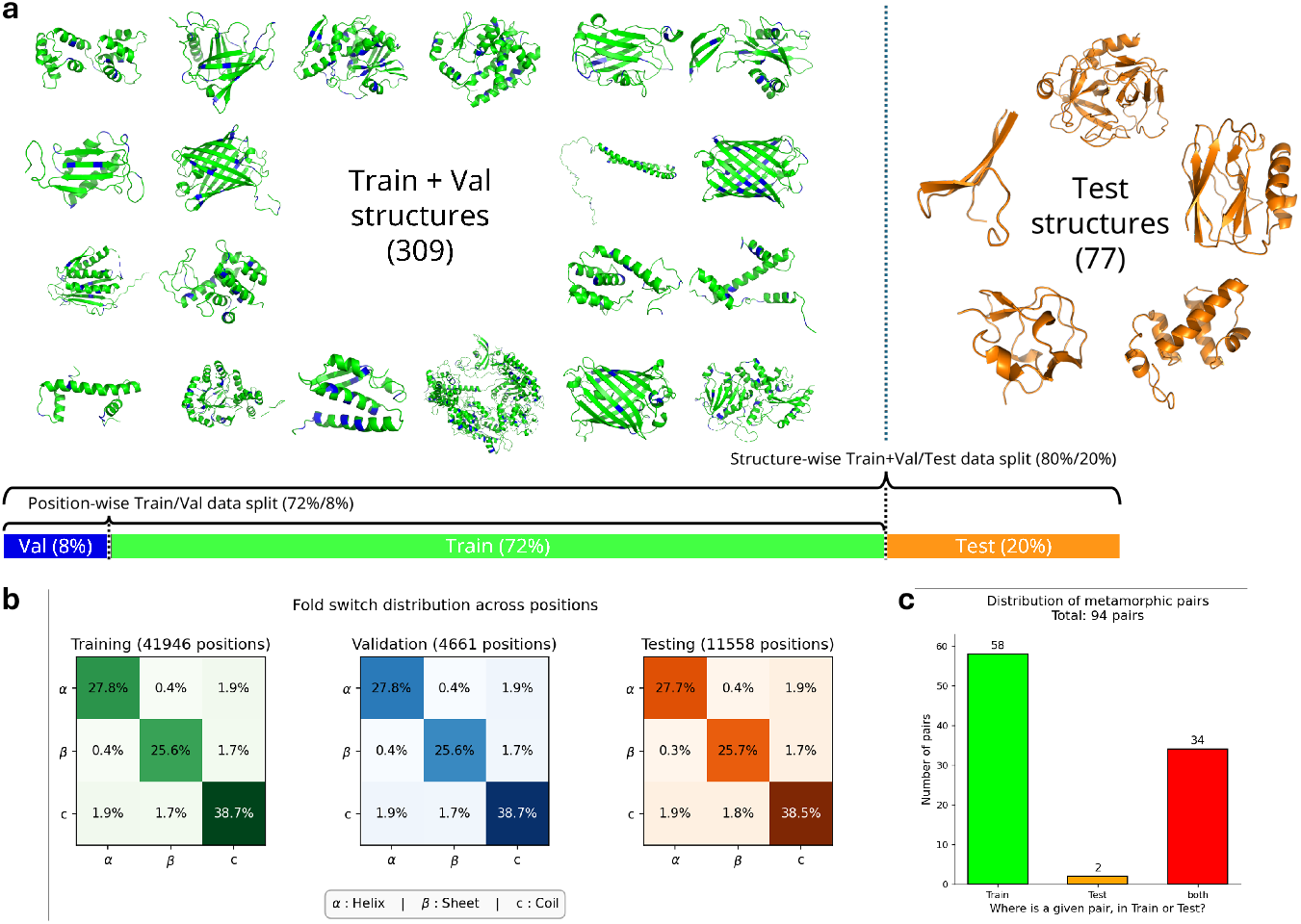
**a**, An illustration of the structure-wise train+val/test data split and the position-wise train/val data split. 3D structures of a representative set of proteins is shown here. **b**, The train/val/test distributions of the 9 possible fold-switches across positions. **c**, The distribution of the metamorphic pairs between train and test data.

### 4.2 Feature engineering for residue-wise structural context

We discuss below the different feature engineering (*ℱ*) that we use to create the feature vectors *xi*, which encode the structural context of a residue at a given position *i* in a protein. We explore, broadly, two categories of feature engineerings, handcrafted and protein language model (PLM) based automated feature engineerings.

#### *k*-Neighbour Contact Separation Interval Composition (kNCSIC) feature

We introduce here a novel handcrafted feature engineering to encode the structural context of a residue at a given position in a protein. We now discuss its construction. It is similar to the CSIC features proposed in [13]. First, we create a contact map *C* for a given input structure. A contact between two residues is defined in the contact maps if the distance between the centre of mass of the two residues is less than 7Å (angstroms)[1]. The contacts of residue are then divided into 4 categories based on the sequence separation of the contacts: (1) neighbourhood contacts, 2-5 residues apart; (2) short-range contacts, 6-11 residues apart; (3) mid-range contacts, 12-23 residues apart; and (4) long-range contacts, more than 24 residues apart [1]. For a given sequence position *i* from a PDB structure, we encode the following from the (2*k* + 1)-mer centred at the *i*^*th*^ residue:

- We encode the (2*k* +1)-mer by concatenating 2*k* +1 one-hot encoded vectors each of 20 dimensions. This results in (2*k*+1) × 20-dimensional feature vector.
- For each of the 2*k* + 1 residues, we look at the amino acid composition frequency counts of their contacts at the 4 contact separation ranges, defined above. This results in a (2*k* + 1) × 4 × 20 dimensional vector.
- For each position, we will also use the secondary structural element (SSE) of the (2*k* + 1)-mer, i.e. *α*/*β*/*c*. This will be a concatenation of (2*k* + 1) one-hot encoded vectors, each being 3-dimensional, resulting in a total of (2*k* + 1) × 3 dimensions.

Thus, overall, we compute a (2*k*+1) × 103-dimensional feature vector as described above. We try *k* values from 1 to 12 to train the classifiers.

#### Protein language model features

In addition to the handcrafted kNCSIC features, we also use pre-trained representations from four state-of-the-art multimodal protein language models (PLMs) that use both sequence and structure information. Specifically, we extract residue-level embeddings from the following structure-aware models:

- **FusionProt**[10] *(4352-dimensional)*: Uses a fusion token as a means of bidirectional fusion between a sequence-based PLM and a graph neural network-based structure encoder.
- **SaProt**[17] *(1280-dimensional)*: Reduces the structure into a sequence of tokens using Foldseek [**?**] and trains them along with amino acid tokens using a traditional PLM.
- **ProtSSN**[19] *(1280-dimensional)*: Uses the residue embeddings of the input sequence as node features of the protein structure-based graph, which is then fed to a graph neural network-based encoder.
- **ProSST**[12] *(768-dimensional)*: Processes a protein’s sequence and 3D structure through a two-step pipeline that converts 3D protein structure into a discrete structure token sequence and combines them with amino acid sequences using disentangled attention in a transformer-like model.

For a given input protein, we process the structural and sequence data through each respective pre-trained PLM and extract a fixed-dimensional dense embedding vector for every sequence position.

### 4.3 Design of classifiers

For each of the feature engineering described in Section 4.2, we train Support Vector Machine (SVM) classifiers to predict whether a given position in a protein structure switches fold or not. We evaluate linear, polynomial (degrees 2 and 3), and radial basis function (RBF) kernels. The classifiers are trained by minimising a weighted hinge loss. The validation set is used to tune the SVM regularisation parameter *C* and kernel-specific parameters. See appendix Section B for additional details on hyperparameter tuning. The best parameter is tuned based on the F1-score on the validation set, calculated from the precision and recall of the switch class. The classification performance is also evaluated using this F1-score on the test set.

As formalised in Section 3, we train both binary (switch vs. no-switch) and multi-class transition classifiers (utilising both One-vs-One and One-vs-Rest strategies). Furthermore, we train the classifier in yet two more ways: input-agnostic and input-dependent routing architectures:

- **Input-agnostic approach**: a single, global SVM evaluates all residues.
- **Input-dependent approach**: a routing mechanism assigns residues to one of three specialised SVMs trained exclusively on feature vectors from positions having the same secondary-structure type (*α*-helix, *β*-sheet, or coil). Consequently, the model dynamically applies the most appropriate classifier based on the input residue’s secondary structure.

#### Classifier evaluation

To robustly assess the performance of our models on the highly imbalanced dataset, we primarily rely on the F1-score computed specifically for the positive “switch” class. For the binary classification task, evaluating this metric is straightforward, as the model directly predicts the binary label (*yi* ∈ {0, 1}).

On the other hand, for the granular multi-class task, the model predicts the specific secondary structure state (*yi* ∈ {*α, β, c*}) of the target conformation. A direct comparison of these predicted labels does not inherently distinguish between fold-switching and non-switching events. To evaluate the effective switch F1-score for multi-class models, we dynamically map the predicted structural states back to a binary transition space. We do this by comparing the predicted target state (*yi*) against the residue’s known initial state in the input structure (*si*). A positive switch event is recorded if and only if the predicted state differs from the initial state (*si ≠ yi*); otherwise, it is classified as a no-switch event. This unified mapping ensures that both binary and multi-class models are evaluated on a strictly comparable metric that directly reflects their ability to detect metamorphic behaviour.

## 5 Fold-switch prediction results and error analysis

Below, we discuss the position-wise fold-switch prediction performances of the different classifiers trained using different feature engineerings. Table 1 summarizes the classifier performances.

**Table 1:**
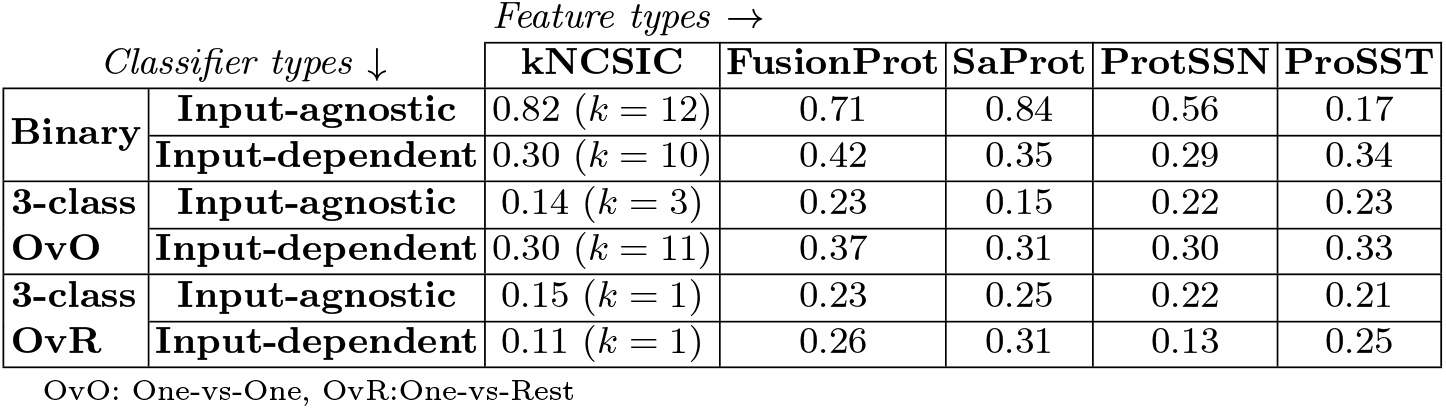
Classification performance summary table. The table reports F1 scores for switch-class prediction on the test data across different classifiers and feature types. For kNCISC features, the scores are reported only for *k* with the highest test score. See classifier evaluation details in Section 4.3.

### Observations from Table 1

We clearly see that classification performance is predominantly higher for the binary input-agnostic classifiers. Under this, the structure-aware PLM SaProt achieves the highest overall performance (F1 = 0.84), closely followed by the handcrafted kNCSIC feature utilising a wide structural window (*k* = 12, F1 = 0.82) and the multimodal FusionProt (F1 = 0.71). On the other hand, the binary input-dependent classifiers and all 3-class classifiers have low test F1 scores (*<* 0.43) across all feature types. Among the 3-class classifiers, the input-dependent One-vs-One classifiers have relatively higher test F1 scores.

### 5.1 Prediction analysis of top performing classifiers

We further analyse the predictions of the top-performing classifiers on the test data. Binary input-agnostic classifiers using SaProt PLM embeddings and kNC-SIC (*k* = 12) are the classifiers with the top-2 test F1 scores. We can see from Table 1 that the SaProt embedding based classifier has only marginally (2%) higher predictive performance than the handcrafted kNCSIC (*k* = 12) feature based classifier. We see from Fig. 2 that the precision and recall for switch-class prediction of the SaProt classifier are also only marginally higher than the kNCSIC classifier. Both classifiers have high precision (≥ 0.97) for switch-class prediction.

**Fig 2:**
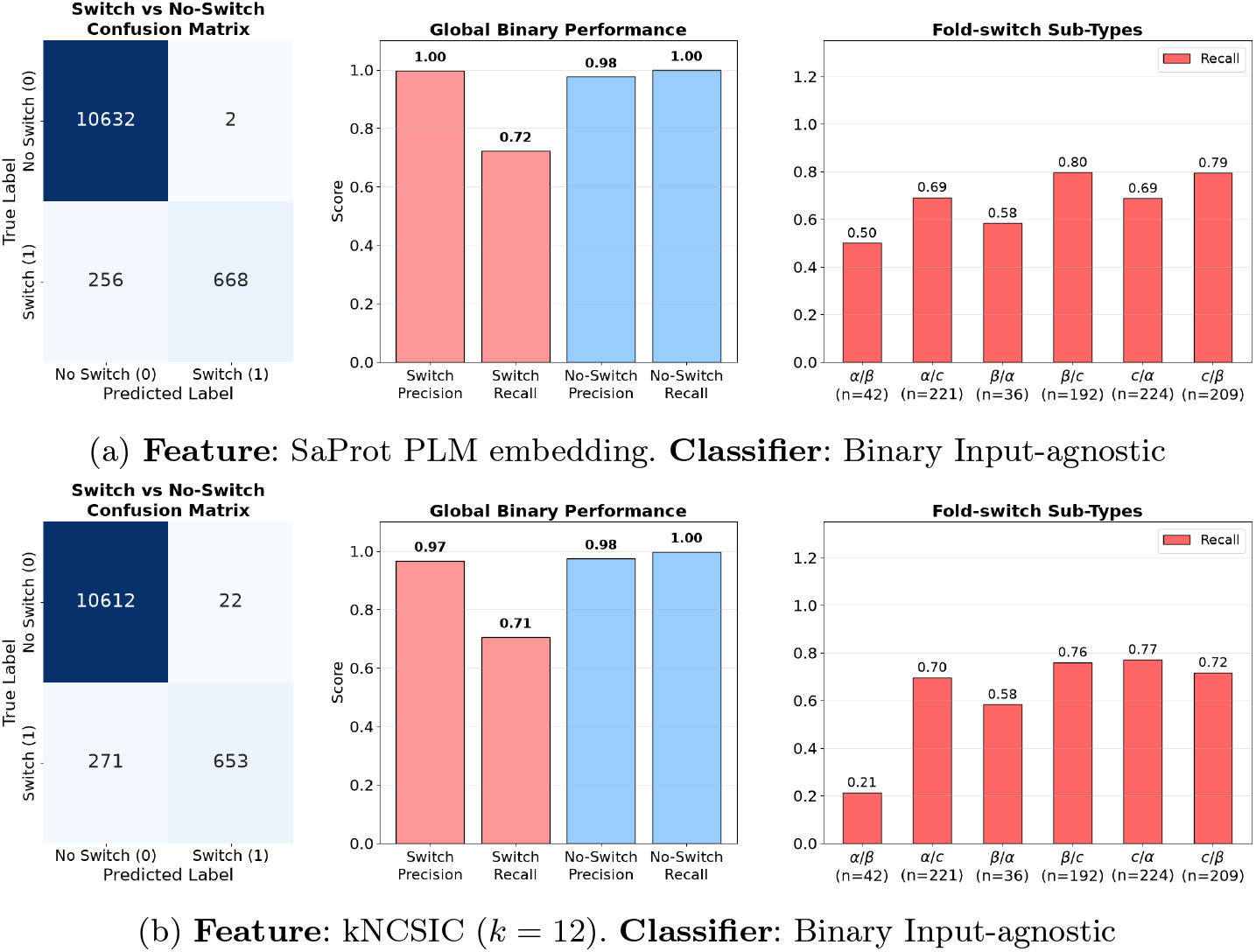
Analysis of the test predictions by classifiers with the highest (top-2) F1 scores from Table 1.

We can see from the rightmost subplots of Fig. 2 that both classifiers have relatively poor performance (*<* 0.59 recall) in predicting *α* → *β* and *β* → *α* switching. The predictive performance is relatively better (*>* 0.68 recall) when the secondary structure of the residue is *c* either in the input structure or in the alternate conformation after fold-switching. The poor performance in the prediction of *α* → *β* / *β* → *α* fold-switching could be due to the low number of samples of this type in our dataset (See Fig. 1b).

### 5.2 Error analysis on metamorphic structures

We performed test error analysis of the classifiers with top-2 F1 scores in Table 1. From the error analysis, we find that the high test F1 score of these classifiers, particularly the PLM based classifier, could be a result of an implicit train/test data leakage in the random 80:20 data split we created for training and testing the classifiers. Below, we discuss implicit train/test data leakage of metamorphic structure pairs and the error analysis of the top-2 classifiers (from Table 1).

#### Implicit train/test data leak for metamorphic structure pairs

In the test set, we had 38 structures that were metamorphic. As illustrated in Fig. 1, 34 of these structures were such that their alternate conformation structure was present in the train set, while 4 structures (i.e. 2 pairs) were there such that both the structural conformations were present in the test set. Thus, the 34 metamorphic test structures with their alternate conformations in the train set exhibit implicit data leakage, as the two conformations share the exact same sequence, differing only in structure due to fold-switching. Below, we present an error analysis of the top-2 classifiers for the two types of metamorphic structures in the test data: 34 structures with train leakage and 4 leak-proof structures.

- **Error analysis of best PLM (SapProt) classifier**. Among the 34 metamorphic test structures with train leakage, the best PLM (Saprot) classifier predicted fold-switching for 33 of them. However, for the 4 leak-proof metamorphic test structures, it predicted fold-switching for only 1 structure. On the 39 monomorphic structures in the test set, it predicted 0 false positives. This suggests a possible sequence memorisation by the PLM classifier for detecting fold-switches.
- **Error analysis of best kNCSIC (***k* = 12**) classifier**. The best kNCSIC (*k* = 12) classifier also predicted fold-switching in 33 of 34 metamorphic test structures with train leakage. Crucially, for the 4 leak-proof metamorphic test structures, it predicted fold-switching in all 4. Furthermore, of the 39 monomorphic structures, it yielded false positives on only two. The fact that the classifier detected fold-switching in completely unseen sequences – while maintaining a low (non-zero) false-positive rate on monomorphic structures – demonstrates that it hasn’t simply memorised sequences.

The above analysis suggests that the PLM-based classifier relies heavily on memorised sequence features from the training set. In contrast, the kNCSIC classifier does not rely exclusively on sequence memory; its superior generalisation on the leak-proof test data indicates that it uses underlying structural features. However, we acknowledge that the size of the strictly leak-proof test data (only 4 structures) is small.

## 6 Discussion

### What do the prediction performances say?

The large performance gap between binary input-agnostic classifiers and the 3-class or input-dependent variants suggests that predicting the *propensity* to switch is much more tractable than explicitly predicting the alternate secondary structure state. Furthermore, the fact that the handcrafted kNCSIC (*k* = 12) feature performed almost on par with the massive, pre-trained SaProt embedding suggests that complex attention mechanisms offered only marginal gains over the local structure encoding handcrafted features.

### Limitations of the work

A limitation of this work is the restriction of the binary input-agnostic formulation. While accurate at predicting *if* a switch will occur, it provides no information on *what* secondary structure the residue will adopt, limiting its direct utility for downstream structural modelling. Another limitation is the classifier’s poor performance in predicting rare *α* ↔ *β* transitions compared to transitions involving coil states (*c*). This could be because large-scale topological rearrangements between rigid secondary structures (*α*-helix and *β*-sheets) are both thermodynamically rarer and severely underrepresented in the dataset.

However, a more serious limitation of the work – something we spotted during the review period – is the implicit data leakage in the train/test split, which calls into question the high predictive performance of the classifiers. As highlighted in Section 5.2, 34 of the 38 metamorphic test structures suffer from implicit sequence leakage into the train set. Thus, the high F1 scores of the best classifiers are due to this data leakage. The classifiers could have memorised the sequence features of the metamorphic structures in the train set, enabling accurate fold-switching predictions for their alternate conformations present in the test set. Even though the handcrafted kNCSIC performed better on the held-out leak-proof metamorphic structure data, the small size of the data (only 4 structures) makes its generalisation ability inconclusive.

To address the leakage, we conducted preliminary experiments using a dataset that includes only a single structural conformation of a metamorphic protein, thereby ensuring no data leakage. Consequently, we are left with a very small sample size of metamorphic structures in the dataset. In these experiments, we find that F1 scores drop significantly, to less than 0.5, for both PLM-based and handcrafted feature-based classifiers. Although PLMs are pre-trained on large amounts of data and have high expressive power, their low predictive performance on this task may stem from the paucity of available metamorphic structure data for training. Thus, more specialised feature engineering and classifier design could be required to achieve good predictive performance in the position-wise fold-switch prediction task. Designing interpretable handcrafted features, such as kNCSIC, could also enable domain-specific insights into the fold-switching phenomena as illustrated in a recent paper [13].

## Acknowledgments

The computational resources used for this study were procured using the contingency grant of the Prime Minister’s Research Fellowship (PMRF) awarded to Pranav Machingal.

## Appendix

### A Details on dataset

We use the dataset of metamorphic and monomorphic proteins from [18], which they themselves collected from [16,7]. The dataset from [18] contained 189 metamorphic structures, out of which the PDB (protein data bank) IDs 4KSO, 1F16, 8E6Y did not have their metamorphic pair structures, i.e. their alternate conformation. We deleted the structures 1F16 and 8E6Y from the dataset, and added 5N8Y which is the alternate conformation of 4KSO. Thus, resulting in 94 pairs of metamorphic proteins, with a pair being two alternate conformations of the same protein.

We do not use all the positions of the PDB structures of the metamorphic proteins for feature extraction. For each metamorphic pair, we first find the longest common contiguous subsequence (*lccs*) from the sequences of the two PDB entries. The feature vectors are extracted only for these positions, which are part of the *lccs*. And thus, for metamorphic pairs, only these feature vectors coming from their *lccs* are used for training and testing of the classifier. For monomorphic proteins, the feature vectors are extracted from all sequence positions.

### B Hyperparameter tuning for SVM

To optimize the standard SVM hyperparameters (regularization parameter *C* and RBF/polynomial kernel parameter *γ*) alongside our custom sample-wise weighting parameter (*a*), we utilize Optuna [2], an automated hyperparameter optimization framework. The tuning is driven by the Tree-structured Parzen Estimator (TPE) algorithm [4], which employs Bayesian optimization to efficiently explore the continuous parameter spaces and dynamically focus on regions yielding the highest validation scores.

For each feature representation and classification architecture, we evaluate four distinct kernel variants: linear, radial basis function (RBF), degree-2 polynomial, and degree-3 polynomial. Each kernel undergoes an independent optimization study consisting of exactly 80 trials.

#### Parameter search spaces

Rather than evaluating a static grid, we define bounded probabilistic continuous search spaces for Optuna to sample from:

- **Regularization and kernel parameters:** Because *C* and *γ* are highly sensitive to orders of magnitude, they are sampled logarithmically.

- *C* ∼ LogUniform(10^*™*6^, 10^3^)
- *γ* ∼ LogUniform(10^*™*6^, 10^2^) for the RBF kernel.
- *γ* ∼ LogUniform(10^*™*6^, 10^*™*1^) for polynomial kernels. (Additionally, the polynomial *coef* 0 parameter is sampled linearly from Uniform(0.0, 1.0)).

- **Sample-wise class weights:** To handle class imbalance, we sample-wise reweigh the Lagrangian multiplier (*αi*) bounds in the SVM dual formulation. The bound *αi* ≤ *C* becomes *αi* ≤ *C* × *ri*. In the SVM primal, this results in the hinge loss *ξi* for sample *i* being reweighted to *riξi* in the objective function. We assign minority (fold-switching) and majority (no-switch) weights using *r*+ and *r™*, governed by a single continuous parameter *a*:

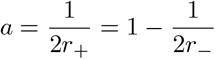

Since *a* must mathematically remain strictly between 0 and 1 to yield valid positive weights, it is sampled continuously across this valid range:

- *a* ∼ Uniform(0.0001, 0.9999)

During each trial, the SVM model is instantiated with the suggested parameters, trained using the derived sample weights, and evaluated. The TPE algorithm dynamically builds probability density functions of the parameter space based on the resulting switch F1-score on the validation set, continuously updating its priors to efficiently steer the search toward the global optimum across the 80 trials. The optimal configuration across all four kernels is then selected as the final model.

